# MicrobiomePrime: A primer pair selection tool for microbial source tracking validated on a comprehensive collection of animal gut and fecal waste microbiomes

**DOI:** 10.1101/2025.07.08.663663

**Authors:** Tanja Zlender, Lucija Brezočnik, Vili Podgorelec, Maja Rupnik

## Abstract

Identifying primer pairs for use in microbial source tracking (MST) can be challenging. The primer pairs must amplify markers with high sensitivity and specificity for a particular microbiota source, while factors such as melting temperature, secondary structure formation, and amplicon length must be carefully considered to ensure efficient amplification. To accomplish this, previous MST studies relied on complex, multi-step analysis, which was labor-intensive and time-consuming. In this paper, we present MicrobiomePrime, a tool for identifying primer pairs for amplification of source-associated markers by analyzing amplicon sequences. Our approach is based on splitting amplicon sequences from target sources into K-mers, which are treated as potential primers in an *in silico* PCR. As proof of concept, we applied the tool to identify markers associated with pig feces, generating thousands of primer pairs. Among nine primer pairs selected for laboratory validation, six primer pairs exhibited *in vitro* specificity higher than 93 %. Among them, four were 100 % specific. While originally developed for MST, MicrobiomePrime has broader applicability, enabling the design of primers targeting any molecular marker based on amplicon sequencing data.

MicrobiomePrime is available on GitHub: https://github.com/tanjazlender/MicrobiomePrime.

**Author summary:** Detecting molecular markers is one of the key approaches for identifying specific traits or differences. One approach for the detection of molecular markers involves amplifying them with primer pairs that are highly specific, sensitive and carefully designed to ensure optimal assay performance. Existing methods for designing these primers are often labor-intensive, involving complex, multi-step analyses. To address this challenge, we developed MicrobiomePrime, a tool that offers a streamlined design of source-associated primer pairs by analyzing sequencing data. It eliminates the need for complex multi-step analysis and enables a rapid development of novel PCR assays for the detection of molecular markers. The method is broadly applicable, however, we have developed it for microbial source tracking sources of fecal contamination. Identifying sources of contamination is namely essential for controlling microbial spread, improving sanitation practices, and ensuring effective remediation of affected environments.

## Introduction

Microbial contamination poses a significant concern across various sectors, impacting public health, food safety, and environmental quality. By understanding contamination pathways, we can enhance infection control in healthcare [1–3] and veterinary settings [4], improve sanitation in the food industry [5–7] and trace sources of fecal contamination in environmental samples, including recreational and drinking water [8–13].

A widely used method for identifying contamination sources is library-independent microbial source tracking (MST). This approach is based on detecting microbial genetic markers that act as unique fingerprints of particular sources of contamination [9, 13]. To identify these markers in polluted samples, they are typically amplified in a PCR reaction, using a highly sensitive and specific pair of primers [6, 13–15].

However, selecting optimal primers for MST can be challenging. Early methods focused on identifying source-associated markers before designing corresponding primer pairs. These studies relied on various techniques including amplification of specific bacterial genes (e.g. the 16S rRNA gene of *Bacteroidales*), followed by differentiation of target and non-target DNA fragments based on competitive hybridization [16, 17], amplicon length, restriction sites [18] and denaturation behavior [19]. Amplicons from target samples that differed from those in non-target samples were then sequenced and used for primer design. In other cases, Sanger sequencing was performed directly after amplification and the sequences of target and non-target samples were compared through computational analysis (Fig 1) [20–23].

**Fig 1.**
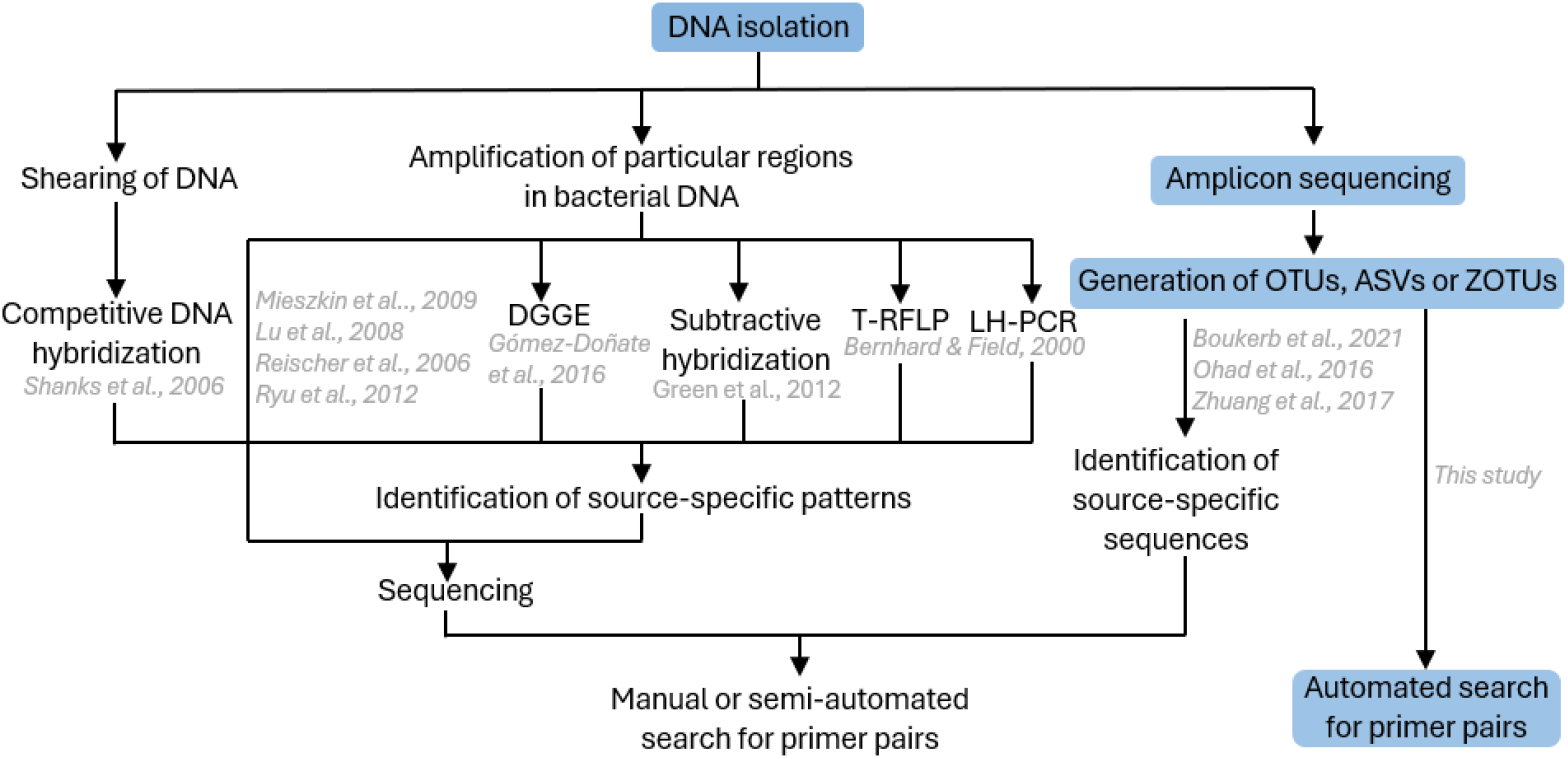
An overview of prior methodologies and our approach for identifying source-associated primer pairs. Steps in our approach are highlighted in blue. DGGE – denaturing gradient gel electrophoresis, T-RFLP – terminal restriction fragment length polymorphism, LH-PCR – length heterogeneity PCR, OTU – operational taxonomic unit, ASV – amplicon sequence variant, ZOTU – zero-radius OTU.

All of the early methods used were very labor-intensive and time-consuming. Furthermore, the process of identifying new MST markers and corresponding primer pairs was manual or semi-automated and included multiple steps such as sequence alignments, clustering, constructing distance trees, performing homology searches in the GenBank database and designing primers, either manually or using specialized software [18, 22, 23].

With the advancements in next-generation sequencing (NGS) technologies, the ability to generate large volumes of data in a short time frame has increased significantly. However, this has introduced new challenges, particularly in the management and analysis of substantially larger datasets. As a result, identifying suitable primer pairs in NGS data remains a complex, multi-step process that still heavily depends on the same methods of sequence analysis used in the earlier approaches [24, 25].

To address these challenges, we developed MicrobiomePrime - a specialized bioinformatic tool that streamlines the identification of primer pairs with high specificity and sensitivity for amplifying markers from target microbiota source(s). As proof of concept, we used MicrobiomePrime to identify primer pairs for the amplification of pig-associated markers from a 16S rRNA sequencing dataset of 767 non-human fecal and fecal waste samples collected in this study, along with 186 human fecal samples obtained from another study and validated them *in vitro*.

## Design and implementation

### The MicrobiomePrime tool

MicrobiomePrime was developed for the analysis of high-throughput amplicon sequencing data, such as variable regions of the 16S rRNA gene, fungal ITS regions and mitochondrial DNA. It is compatible with an x86-64 Linux OS (tested on Ubuntu) and requires the installation of both Python and R. To streamline setup, users are provided with a pre-configured Conda environment, including dependencies such as Biopython and R-based libraries. MicrobiomePrime incorporates OpenMPI for parallelization, optimizing computational efficiency on high-performance systems. This feature is particularly useful for processing large datasets, while smaller datasets can be analyzed without parallelization.

MicrobiomePrime works by splitting target amplicon sequences (sequences found in microbiota of interest) into K-mers of desired primer length. K-mers meeting predefined primer design, sensitivity and specificity criteria are then combined into primer pairs. The primer pairs undergo *in silico* PCR simulations on template sequences from target and non-target sources using ThermoNucleotideBLAST [26]. This step assesses each pair’s sensitivity and specificity. The individual steps of the MicrobiomePrime analysis, including data preprocessing, are shown in Fig 2 and described in detail in the section Detailed steps in MicrobiomePrime analysis.

**Fig 2.**
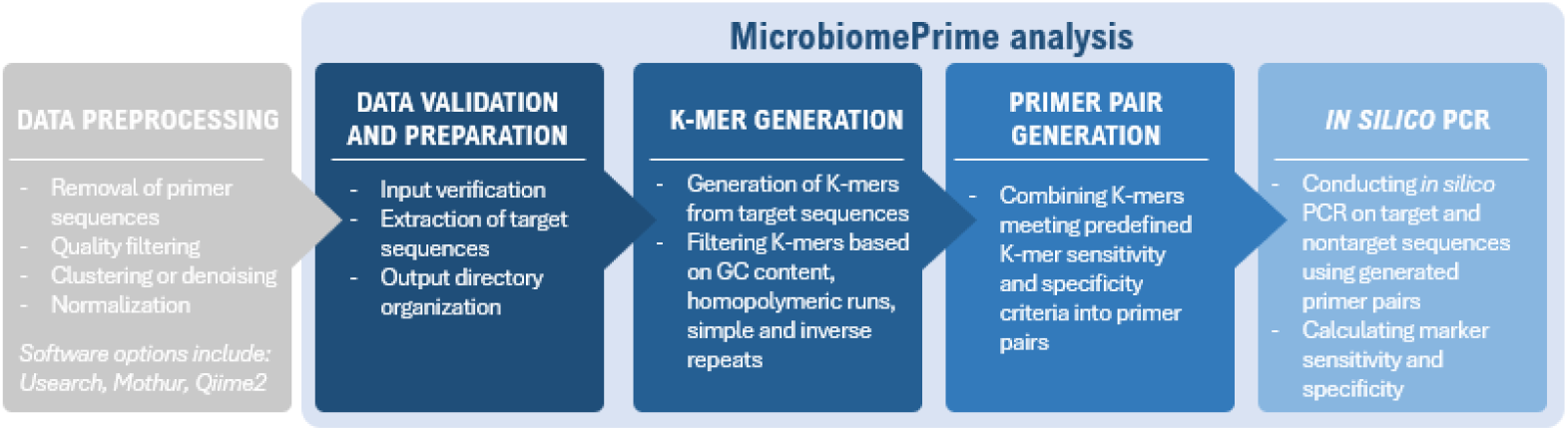
Steps for identifying primer pairs for amplifying source-associated markers using MicrobiomePrime.

### Overview of inputs, parameters and outputs

#### Input files

MicrobiomePrime requires four inputs: a FASTA file containing both target and non-target sequences, a relative abundance table linking sequence IDs to samples, a metadata file, and a taxonomy file. These files can be generated using standard software for amplicon sequence preprocessing [27–31].

#### Parameters and settings

Users can set up to five target sources and specificity exceptions (samples excluded from sensitivity and specificity calculations). This feature is particularly useful for excluding samples of unknown origin from the analysis or when the detection of certain samples is acceptable, although they are not the primary target (e.g. pig manure can be set as an exception when pig feces are the target). Additionally, users can define K-mer size, sensitivity and specificity cutoffs, and PCR amplification parameters, including amplicon length, minimum and maximum primer melting temperature (T_m_), primer delta G, allowable mismatches to the template, and primer clamp. Users can also set the number of CPUs and memory allocated for the task when using a Slurm Workload Manager.

#### Output files

MicrobiomePrime produces a primary results table that summarizes the performance and characteristics of primer pairs, along with supplementary tables containing information about detected sequences. The primary results table includes key metrics such as specificity, sensitivity, and abundance. It also provides taxonomic attribution, amplicon sizes, mismatch counts to the template DNA, and important primer pair features, such as melting temperatures of individual primers, hairpins, homodimers, and heterodimers. Additionally, the table offers details on amplification results for samples marked as exceptions. Example of a primary output table is shown in S2 Table.

### Detailed steps in MicrobiomePrime analysis

The following section provides a detailed explanation of the steps involved in the MicrobiomePrime analysis shown in Fig 2.

#### Data preprocessing

Prior to MicrobiomePrime analysis, raw amplicon sequences need to be preprocessed. This involves the removal of primer sequences, quality filtering, chimera detection and removal, sequence clustering or denoising and normalization. Each sequence is assigned a unique Sequence ID (SeqID) in the format of letters followed by numbers (e.g., Otu001, Otu002). Recommended preprocessing tools include Usearch [27, 28], Qiime2 [29], DADA2 [31] and Mothur [30]. The sequence ID table (e.g., OTU table) should be normalized, and the values should be recalculated as relative abundances, where the sum of abundances for each sample equals 1.

#### Data validation and preparation

This step ensures that all input files are formatted according to our requirements and creates and organizes directories for storing analysis outputs. It then creates a list of sequence IDs found in target samples and extracts corresponding sequences from a FASTA file which are used for K-mer generation in the next step.

#### K-mer generation

MicrobiomePrime segments sequences from target sources (e.g., pig feces) into smaller units, known as K-mers, which are the length of desired PCR primers. This segmentation is achieved using a sliding window approach with a one-base pair shift, as illustrated in Fig 3. K-mers are excluded from the analysis if they contain homopolymeric runs (four or more identical nucleotides in a row), simple repeats (repeated sequences of four or more nucleotides), inverse repeats (self-complementary motifs of four or more nucleotides) or have a GC content outside the range of 45 % to 60 %. These exclusions are necessary to ensure that the K-mers, which will later be used as PCR primers, do not introduce amplification errors, or form secondary structures that could impair primer performance in PCR.

**Fig 3.**
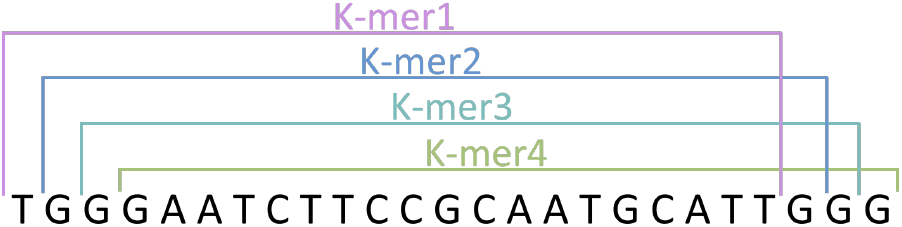
Generation of K-mers. The K-mers were generated by employing a sliding window approach with a one base pair shift.

#### Primer pair generation

Primer pairs are esentially pairs of K-mers meeting our sensitivity and specificity criteria: both primers in a pair must meet the preset sensitivity criteria, while only one must meet the specificity criteria. Sensitivity and specificity of each K-mer are calculated using Eq (1) and Eq (2) respectively, where TP is the number of target samples in which the K-mer was detected, FN is the number of target samples in which the K-mer was not detected, TN is the number of non-target samples in which the K-mer was not detected and FP is the number of non-target samples in which the K-mer was detected.

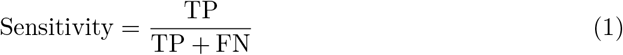

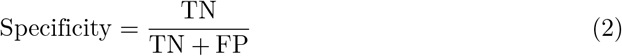

#### *In silico* PCR analysis

In the final step, the generated primer pairs are are subjected to *in silico* PCR simulations using template sequences from both target and non-target sources. The PCR is conducted using ThermoNucleotideBLAST [26].

Sensitivity and specificity of each primer pair are calculated as described earlier (using Eq (1) and Eq (2)), with different definitions of the components: TP is the number of target samples amplified using the primer pair, FN is the number of target samples not amplified by the primer pair, TN is the number of nontarget samples not amplified by the primer pair and FP is the number of nontarget samples amplified by the primer pair. Primer pairs that demonstrate optimal *in silico* performance are considered candidate MST primer pairs.

### Implementation

We used MicrobiomePrime to identify pig-associated primer pairs from a 16s rRNA gene dataset comprising of 715 non-human fecal samples, 186 human fecal samples and 52 animal waste samples. The specificity of these primer pairs was validated *in vitro*, demonstrating their suitability for MST applications.

#### Sample and data acquisition

To ensure a diverse dataset, we incorporated both newly acquired data from our laboratory work and publicly available data. We collected fecal samples from various non-human mammalian and avian sources and animal waste (manure heap, manure heap leachate, slurry pit and slurry tank samples) across Slovenia from December 2020 to September 2023. In total, we gathered 515 fecal samples from mammals, 200 from birds, and 52 animal waste samples derived from cattle and pigs. DNA was isolated from these samples, which then underwent amplicon sequencing targeting the V3-V4 hypervariable regions of the 16S rRNA gene. Detailed protocols for sampling, DNA isolation and amplicon sequencing is provided in S1 Appendix. The sequences are deposited in the NCBI SRA (Accession number PRJNA1191222). Additionally, 186 human fecal sequences were obtained from another study [32]. All samples included in the study are listed in S1 Table along with their taxa.

#### Data preprocessing and filtering

Raw sequencing data was processed using Usearch version 11.0.667 [27, 28]. First, we merged paired-end raw reads and removed reads with more than one expected error. Unique sequences were denoised using UNOISE algorithm, resulting in the formation of zero-radius operational taxonomic units (ZOTUs). We then removed sequences shorter than 400 bp from the analysis and assigned taxonomy to the remaining sequences using the RDP training set (v.18) with a bootstrap threshold value of 0.8.

Using R v.4.1.3, we excluded non-bacterial ZOTUs, ZOTUs classified as chloroplasts at the order level and samples unidentified on the genus level from the ZOTU table. The ZOTU table was then rarefied to 10,000 reads per sample using the Vegan package v.2.6.6.1 and samples with fewer than 10,000 reads were removed from the analysis. Rare ZOTUs, defined as those present in 30 % or fewer samples within each microbiota source and with a relative abundance of 0.01 % or less, were also excluded. The final step of data preprocessing involved calculating relative abundances in the filtered ZOTU table.

#### MicrobiomePrime analysis

For MicrobiomePrime analysis, we selected pig feces as the target for primer design, utilizing a K-mer length of 22. Pig waste samples were excluded from sensitivity and specificity calculations, as the presence of bacteria from pig feces is expected in pig waste and considered acceptable. The defined variables for the analysis are listed in Table 1. The analysis was performed on a high-performance (HPC) computing cluster utilizing 200 CPUs and 150 GB of RAM, managed by the Slurm job scheduling system.

**Table 1.** Variables used in test MicrobiomePrime analysis.

| Variable | Setting/Threshold |
| --- | --- |
| target | Pig |
| specificity_exception | Pig waste |
| kmer_size | 22 |
| kmer_sensitivity_cutoff | 80 |
| kmer_specificity_cutoff | 98 |
| marker_sensitivity_cutoff | 80 |
| marker_specificity_cutoff | 98 |
| min_amplicon_length | 75 |
| max_mismatch | 2 |
| primer_clamp | 2 |

#### The selection of primer pairs

In the output table of MicrobiomePrime, we filtered out primer pairs with forward and reverse primer melting temperature differences greater than 3 °C. From the remaining primer pairs, we selected nine that represented diverse taxa and relative abundances. We prioritized those that satisfied empirical primer design rules (marked as PCR_VALID in the output table) and exhibited low melting temperatures for hairpins, homodimers, and heterodimers.

#### *In vitro* validation of primer pairs

We validated the selected primers (IDT, Coralville, IA) using end-point PCR on composite DNA samples from 18 sources: cattle, pig, horse, sheep, goat, chicken, dog, cat, roe deer, red deer, fallow deer, pigeon, swan, mallard, nutria, wild boar feces, as well as cattle and pig waste. The DNA concentration of five random samples from each source was first measured using a Qubit Flex Fluorometer (Invitrogen), then diluted to 1 ng/μL and combined to form composite samples.

In the first PCR experiment, we tested primer pairs on the target composite sample (pig fecal DNA) at four different annealing temperatures. The optimal annealing temperature was determined based on the strongest band observed after electrophoresis; if multiple bands showed similar intensity, we selected the highest annealing temperature for further experiments (Table A in S1 Appendix). In the second experiment, we evaluated whether the markers amplified non-target samples using the previously selected annealing temperatures.

All PCR reactions were performed in 20 μL mixtures containing PCR buffer (Roche), 0.2 μM of each primer, 200 μM PCR nucleotide mix (Roche), 0.02 U of Taq polymerase (Roche) and 50 pg/μL DNA. The PCR cycling conditions were as follows: an initial denaturation at 95 °C for 3 min; 30 cycles of 95 °C for 30 s, an appropriate annealing temperature (Table A in S1 Appendix) for 30 s, and 72 °C for 30 s; followed by a final extension at 72 °C for 3 min. The assays were performed on a Biometra TAdvanced Thermal Cycler (Analytik Jena) and included a positive and a negative control.

### Limitations

One key constraint of MicrobiomePrime is that it currently allows users to specify only a single K-mer size (corresponding to primer length) for the analysis, which restricts the ability to combine primers of varying lengths into primer pairs. Although users can manually modify primer lengths in MicrobiomePrime output table by adding or removing nucleotides, this should be approached with caution, as such changes may compromise sensitivity and specificity. Another limitation is that analyzing large datasets can be resource-intensive, requiring substantial memory and CPU usage.

### Ethics statement

Publicly available sequence data from previous study [32] were used, that was approved by Republic of Slovenia National Medical Ethics Committee under approval number [0120-142/2016-2 KME 81/03/16].

## Results

### *In silico* PCR analysis

The MicrobiomePrime analysis was completed over a span of two days and identified 57,484 pig-associated primer pairs that met our *in silico* sensitivity, specificity, amplicon size and primer design criteria. Following an additional filtering step to eliminate primer pairs with melting temperature differences greater than 3 °C, we got 28,041 candidate primer pairs (S2 Table). Among these, nine primer pairs were selected for validation using *in vitro* end-point PCR (Table 2; for selection criteria see section The selection of primer pairs).

**Table 2.**
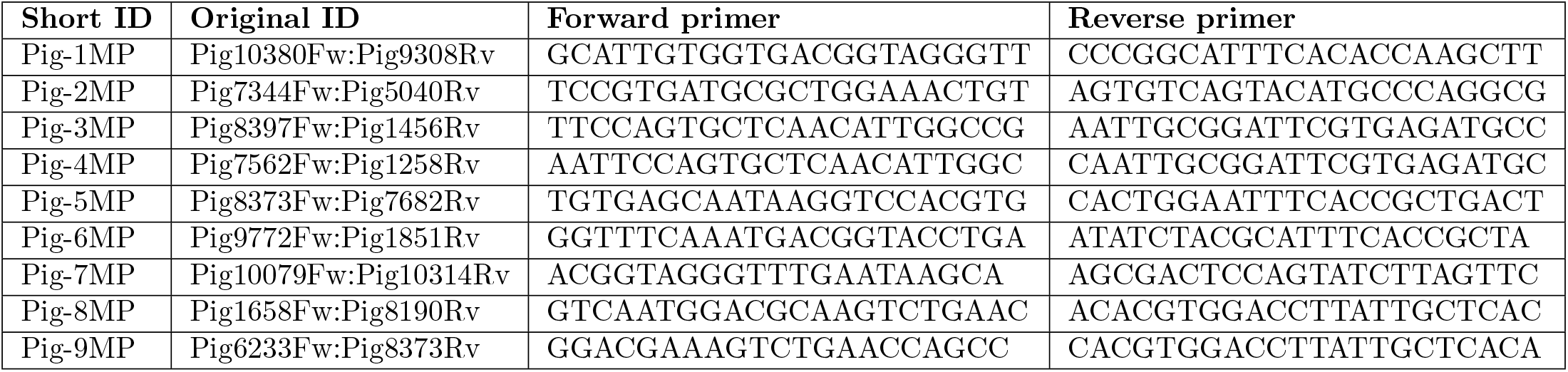
The selected primer pairs with corresponding forward and reverse primer sequences.

### *In vitro* validation

All of the tested primer pairs amplified markers of expected size in target samples using end-point PCR (Table A in S1 Appendix). Our results demonstrate that MicrobiomePrime analysis can effectively screen for highly specific primers, as six out of nine tested primer pairs (Pig-1MP, Pig-2MP, Pig-4MP, Pig-7MP, Pig-8MP, and Pig-9MP) achieved *in vitro* specificities above 93 %, making them promising candidates for MST applications. Pig-3MP also performed well, with 87.5 % specificity, showing limited cross-reactivity only with the horse and nutria samples. However, Pig-5MP and Pig-6MP displayed broad cross-reactivity, amplifying multiple nontarget samples, including cattle, horse, sheep, goat, chicken, deer, and nutria (Table 3).

**Table 3.** *In silico* and *in vitro* PCR results.

| Primer pair | Specificity |  | Detected in pig manure |  | Detected nontarget samples |  |
| --- | --- | --- | --- | --- | --- | --- |
|  | *in silico* | *in vitro* | *in silico* | *in vitro* | *in silico* | *in vitro* |
| Pig-1MP | 100 % (764/764) | 100 % (16/16) | Yes (1/21) | Yes | / | / |
| Pig-2MP | 100 % (764/764) | 93.8 % (15/16) | No | No | / | Horse |
| Pig-3MP | 100 % (764/764) | 87.5 % (14/16) | Yes (1/21) | No | / | Horse, nutria |
| Pig-4MP | 100 % (764/764) | 100 % (16/16) | Yes (1/21) | No | / | / |
| Pig-5MP | 100 % (764/764) | 37.5 % (6/16) | Yes (1/21) | No | / | Cattle, horse, sheep, goat, chicken, roe deer, red deer, fallow deer, mallard, nutria |
| Pig-6MP | 100 % (764/764) | 0 % (0/16) | Yes (3/21) | Yes | / | Cattle, horse, sheep, goat, chicken, dog, cat, roe deer, red deer, fallow deer, pigeon, swan, mallard, nutria, wild boar, cattle manure |
| Pig-7MP | 100 % (764/764) | 100 % (16/16) | No | No | / | / |
| Pig-8MP | 99.6 % (761/764) | 93.8 % (15/16) | Yes (1/21) | No | Horse | Horse |
| Pig-9MP | 99.1 % (757/764) | 100 % (16/16) | Yes (1/21) | Yes | Bear, horse | / |

The numbers in parentheses show the number of positive samples followed by the total number of samples.

The results of specificity testing underscore the importance of validating primers *in vitro* to confirm specificity. While our *in vitro* screening was based on composite samples of 19 microbiota sources, more accurate specificity assessments and sensitivity evaluations will require testing on individual fecal samples.

## Availability and future directions

MicrobiomePrime is available on GitHub: https://github.com/tanjazlender/MicrobiomePrime. The tool has significant potential for further development. Code optimizations could improve its performance, increasing speed and reducing resource consumption, while modifications could expand its functionality. For example, future updates could facilitate the integration of primer pairs with varying lengths, allowing for greater flexibility in primer design. Furthermore, the code could be adapted to identify primers that amplify markers exhibiting differential abundance between target and non-target groups, rather than focusing solely on presence-absence. This enhancement could be particularly valuable for diagnosing gut-related diseases, where specific taxa often show characteristic differences in abundance [33–35]. Finally, the underlying principles of the code could also be adapted for use with shotgun sequencing data.

## Supporting information

**S1 Table. Sample and host taxa information**. This table lists all samples included in the study, along with their associated host taxa and the classification of host taxa into broader groups, which were used as attributes in the MicrobiomePrime analysis.

**S2 Table. MicrobiomePrime output table**. The primary MicrobiomePrime output table for the test target host (pig) containing sensitivity, specificity and relative abundance information along with taxonomic attribution, amplicon sizes, mismatch counts to the template DNA, and important primer pair features, such as melting temperatures of individual primers, hairpins, homodimers, and heterodimers.

**S1 Appendix. Supplementary analysis and method details**. This appendix includes detailed steps of sample collection, DNA isolation, NGS sequencing, and selection of annealing temperatures for *in vitro* validation.

## Notes

### Competing Interest Statement

The authors have declared no competing interest.

### Funding

- Slovenian Research Agency: P3-0387
- European Centre for Disease Prevention and Control: Grant/2021/PHF/23776 (EU/ECDC)

