## Supplementary material for "MicrobiomePrime: A primer pair selection tool for microbial source tracking validated on a comprehensive collection of animal gut and fecal waste microbiomes": S1 Appendix

### Sampling

Fecal samples from various animal sources, including mammals, birds, and animal waste (manure heap, manure heap leachate, slurry pit and slurry tank samples) were collected across Slovenia from December 2020 to September 2023. In total, we gathered 515 fecal samples from mammals, 200 from birds, and 52 animal waste samples derived from cattle and pigs.

Sterile stool containers and sterile centrifuge tubes were used to sample stool samples and animal waste respectively. The samples were promptly placed in ice-filled coolers upon collection and transported to the laboratory within 24 hours. If immediate transportation was not feasible, they were frozen at -20 °C and then transported to the laboratory on ice as soon as possible. Upon arrival, 0.25 g of each fecal sample and solid animal waste sample was frozen at -80 °C until further processing. Liquid animal waste samples (2–4 ml, depending on liquid content) were centrifuged at 10,000 × g for 15 minutes, and the resulting pellet was frozen at -80°C.

### DNA isolation and amplicon sequencing

Total DNA was isolated from fecal samples using the QIAamp Fast Stool Mini Kit (Qiagen, Hilden, Germany) with mechanical disruption (MagNA Lyser; 7000 rpm for 70s). DNA from 0.25 g solid and centrifuged liquid animal waste samples was isolated using the QIAamp PowerFecal Pro DNA Kit (Qiagen, Hilden, Germany). The isolated DNA was stored at -80 °C until library preparation. We sequenced the V3-V4 hypervariable region of the 16S rRNA gene. The region of interest was amplified using primers Bakt_341F (5′-CCTACGGGNGGCWGCAG-3′) and Bakt_805R (5'-GACTACHVGGGTATCTAATCC-3')[1,2]. Library preparation was performed according to the Illumina 16S Metagenomic Sequencing Library Preparation protocol (Illumina, CA, USA) using either KAPA HiFi

HotStart ReadyMix (Kapa Biosystems, MA, USA) or Q5 High-Fidelity DNA Polymerase (New England Biolabs, USA). For samples AF001 to AF372 and CM001 to CM005, KAPA HiFi was used for library preparation, and sequencing was performed on the Illumina MiSeq platform using the MiSeq Reagent Kit V3 (600 cycles). For all other samples, sequencing was performed on the NextSeq 2000 platform using the NextSeq 1000/2000 Reagents (600 cycles). A 10% Phix standard was included in each sequencing run for quality control.

### *In vitro* validation of primer pairs on target samples

The primer pairs were first tested on composite target samples at different annealing temperatures listed in Table A. The annealing temperatures were set based on the mean melting temperatures ($T_m$) of the forward and reverse primer.

**Table A: Melting temperatures and annealing temperatures**. Melting temperatures of selected primers, mean melting temperature of forward and reverse primers and annealing temperatures at which the primer pairs were tested in a PCR reaction.

| **Short ID** | **Original ID** | **Forward $T_m$** | **Reverse $T_m$** | **Mean $T_m$** | **Annealing temperatures (°C)** |
|---|---|---|---|---|---|
| Pig-1MP | Pig10380Fw:Pig9308Rv | 59.8 | 60.8 | 60.3 | 56.6, 58.4, 60.3*, 62.3 |
| Pig-2MP | Pig7344Fw:Pig5040Rv | 61.5 | 61.6 | 61.55 | 58.4, 60.3, 62.3*, 63.7 |
| Pig-3MP | Pig8397Fw:Pig1456Rv | 60.4 | 58.2 | 59.3 | 54.7, 56.6, 58.4, 60.3* |
| Pig-4MP | Pig7562Fw:Pig1258Rv | 56.4 | 57 | 56.7 | 52.7, 54.7, 56.6, 58.4* |
| Pig-5MP | Pig8373Fw:Pig7682Rv | 57.3 | 57.6 | 57.45 | 52.7, 54.7, 56.6, 58.4* |
| Pig-6MP | Pig9772Fw:Pig1851Rv | 55 | 53.8 | 54.4 | 51.3, 52.7, 54.7, 56.6* |
| Pig-7MP | Pig10079Fw:Pig10314Rv | 54.4 | 53.6 | 54 | 51.3, 52.7, 54.7*, 56.6 |
| Pig-8MP | Pig1658Fw:Pig8190Rv | 56.4 | 57.3 | 56.85 | 52.7, 54.7, 56.6, 58.4* |
| Pig-9MP | Pig6233Fw:Pig8373Rv | 59.4 | 57.3 | 58.35 | 54.7, 56.6, 58.4, 60.3* |

The melting temperatures of ordered primers listed here may differ from those in the MicrobiomePrime results table (S2 Table). Legend: *Annealing temperature selected for specificity testing.
